# Genome-Wide Association Study of Body Fat Distribution identifies Novel Adiposity Loci and Sex-Specific Genetic Effects

**DOI:** 10.1101/207498

**Authors:** Mathias Rask-Andersen, Torgny Karlsson, Weronica E Ek, Åsa Johansson

## Abstract

Body mass and body fat composition are of clinical interest due to their links to cardiovascular- and metabolic diseases. Fat stored in the trunk has been suggested as more pathogenic compared to fat stored in other compartments of the body. In this study, we performed genome-wide association studies (GWAS) for the proportion of body fat distributed to the arms, legs and trunk estimated from segmental bio-electrical impedance analysis (sBIA) for 362,499 individuals from the UK Biobank. A total of 97 loci, were identified to be associated with body fat distribution, 40 of which have not previously been associated with an anthropometric trait. A high degree of sex-heterogeneity was observed and associations were primarily observed in females, particularly for distribution of fat to the legs or trunk. Our findings also implicate that body fat distribution in females involves mesenchyme derived tissues and cell types, female endocrine tissues a well as several enzymatically active members of the ADAMTS family of metalloproteinases, which are involved in extracellular matrix maintenance and remodeling.

---

Overweight (body mass index [BMI] >25) and obesity (BMI>30) have reached epidemic proportions globally^1^. Almost 40% of the world’s population are now overweight^2^ and 10.8% are obese^3^. Obesity is set to become the world’s leading preventable risk factor for disease and early death due to the increased risks of developing type 2 diabetes, cardiovascular disease, and cancer^4^.

The distribution of adipose tissue to discrete compartments within the human body is associated with differential risk for development of cardiovascular and metabolic disease^5^. Body fat distribution of fat is also well known to differ between sexes. After puberty, women accumulate fat in the trunk and limbs to a proportionally greater extent compared to other parts of the body, while men accumulate a greater extent of fat in the trunk^6^. Accumulation of adipose tissue around the viscera, the internal organs of the body, has been shown to be associated with increased risk of disease in both men and women^7^. In contrast, the preferential accumulation of adipose tissue in the lower extremities, i.e. the hips and legs, has been suggested as a factor contributing to the lower incidence of myocardial infarction and coronary death observed in women during middle age^8^. The differential distribution of body fat between sexes has been attributed to downstream effects of sex hormone secretion^5^. However, the biological mechanisms that underlie body fat distribution have not been fully elucidated.

BMI is commonly used as a proxy measurement of body adiposity in epidemiological studies and in clinical practice. However, BMI is unable to discriminate between adipose and lean mass, and between fat stored in different compartments of the body. Other proxies that better represent distribution of body fat have also been utilized, such as the waist-to-hip ratio (WHR), waist circumference (WC), and hip circumference (HC). Through genome-wide association studies (GWAS), researchers have identified hundreds of loci to be associated with proximal measurements of body mass and body fat distribution such as BMI^9^, WHR^10,11^ and hip-, and waist circumference^11^. Sex-stratified analyses have revealed sexual dimorphic effects at twenty WHR-associated loci and 19 of these loci displayed stronger effects in women^12^. Body fat mass has also been studied in GWAS by using bio-electrical impedance analysis (BIA) and dual energy X-ray absorptiometry (DXA)^13,14^. BIA measures the electrical impedance through the human body, which can be used to calculate an estimate of the total amount of adipose tissue. The ‘gold standard’ method for measurements of body fat distribution is computed tomography (CT) or magnetic resonance imaging (MRI). However, these methods are costly. A GWAS has been performed for subcutaneous- and visceral adiposity, measured with computed tomography scans, albeit in a relatively limited number of individuals (N=10,577)^15^.

Developments in BIA technology has now allowed for cost-efficient segmental body composition scans that estimate of the fat content of the trunk, arms and legs^16^ (Figure 1a). In this study, we used segmental BIA data on 362,499 participants of the UK Biobank to study the genetic determinants of body fat distribution to the trunk, arms and legs. For this purpose, we performed GWAS on the proportion of body fat distributed to these compartments. We also performed sex-stratified analyses to determine sex-specific effects and performed gene-sex interaction analyses to identify effects that differ between men and women.

**Figure 1.**
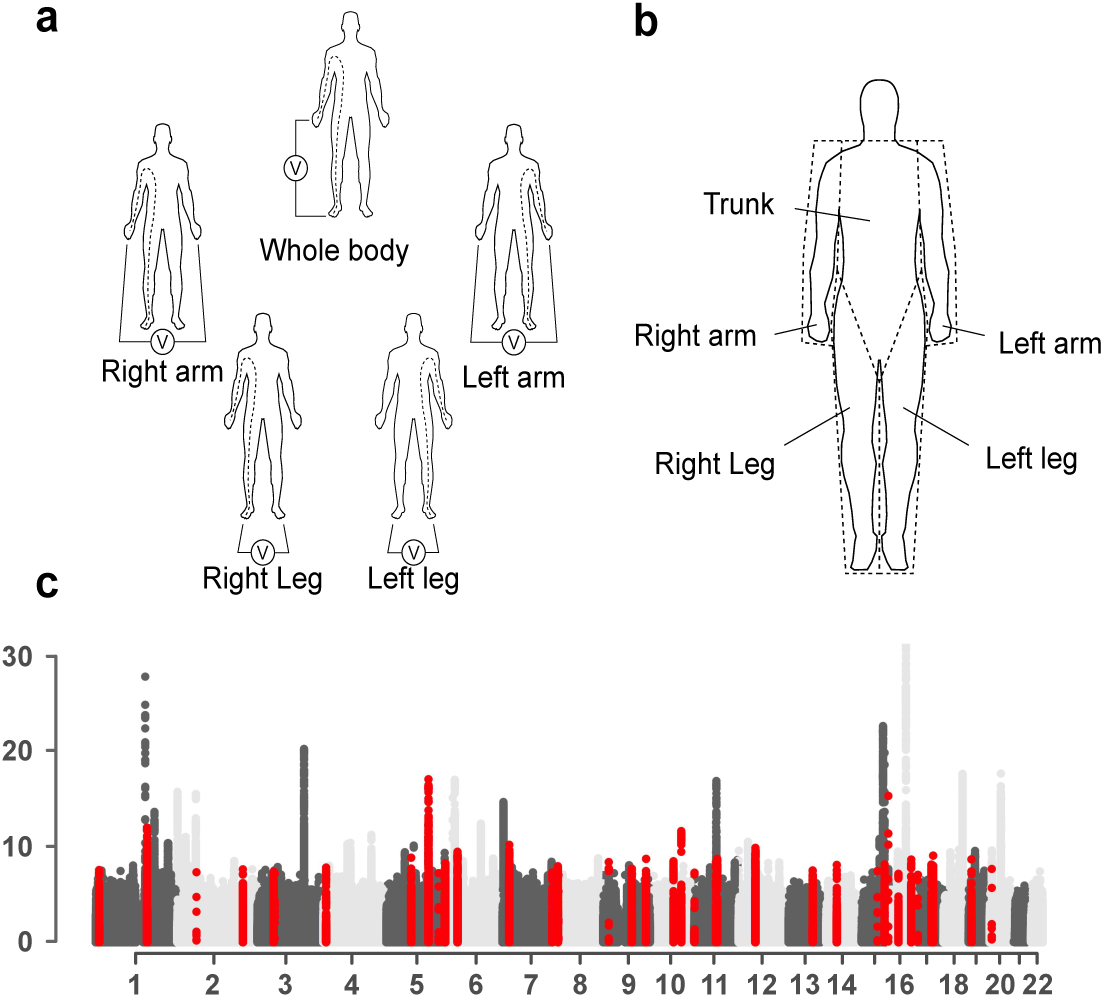
Segmental body impedance analyses. use bioelectrical impedance to estimate body composition: fat mass, muscle mass, etc. Adipose tissue mass, in this study, had been estimated using the Tanita BC-418MA body composition analyzer (**a**). This machine uses an eight-electrode method, which allows for five measurements of impedance. Current is supplied to the front of both feet and the fingertips of both hands. Voltage is measured on either heel or thenar portion of the palms. Body composition is derived from a regression formula for each body part. The formula is derived from regression analysis using height, weight, age and impedance for each body part as predictors for composition of each body part as assessed by DXA (**b**). GWAS for AFR, LFR and TFR were conducted in the UK Biobank cohort and revealed associations with loci that have not previously been associated with standard anthropometric traits. (c) shows a manhattan plot with combined results for association studies of body fat ratios in combined and sex-stratified analyses. Novel loci are highlighted in red.

## Results

The proportions of body fat distributed to the arms – arm fat ratio (AFR), the legs – leg fat ratio (LFR) and the trunk – trunk fat ratio (TFR) were calculated by dividing the fat mass per compartment with the total body fat mass for each participant (Figure 1a). We conducted a two-stage GWAS using data from the interim release of genotype data in UK Biobank as a discovery cohort. Another set of participants, for which genotype data were made available as part of the second release, was used for replication. After removing non-Caucasians, genetic outliers and related individuals, 116,138 and 246,360 participants remained in the discovery and replication cohorts, respectively. Basic characteristics of the discovery and replication cohorts are presented in supplementary Table 1. Women were found to have higher total sBIA-estimated fat mass compared to men in both the discovery and replication cohort, as well as higher amount of fat in the arms and legs. Males had higher average proportion of body fat located in the trunk compared to females (62.2% vs. 50.3%) and women had a larger proportion of body fat located in the legs (39.7% vs. 28.1%). While the total amount of adipose tissue in the arms was estimated to be higher in women compared to men, the fraction of adipose tissue distributed to the arms were similar. Several smaller differences between the discovery and replication cohorts were present (supplementary Table 1), such as some slight differences in height and age between men and women in the discovery and replication cohorts. These differences most likely represent the 50,000 participants for the UK Biobank Lung Exome Variant Evaluation (UK BiLEVE) project that were included in the first release of genotyping data for ~150,000 participants, which were used as a discovery cohort in this study. Selection for UK BiLEVE was conducted with specific consideration to lung function which may reflect the differences in baseline characteristics for this subset of the cohort. However, these differences are unlikely to affect the results from our analyses.

### Genome wide association study for body fat ratio

GWAS was performed for each of the three phenotypes (AFR, LFR and TFR) in the whole discovery cohort (sex-combined) and when stratifying by sex (males and females), adjusting for covariates as described in the method section. A total of 25,472,837 imputed SNPs, with MAF of at least 0.0001, were analyzed in the discovery GWAS. LD score regression intercepts^17^ ranged from 1.00 to 1.03 (supplementary Figure 1, supplementary Table 2), and were used to adjust for genomic inflation. We used the -clump function in PLINK^18^, in combination with conditioning on the most significant SNP, to identify associations that were independent within each GWAS as well as between GWAS for the three body fat ratios (AFR, TFR, or LFR) or strata (males, females or sex-combined) (see methods). In total, 133 independent associations at 114 loci were observed in the discovery analyses (Figure 1c, Supplementary Figure 2-4, Supplementary Data 1). For each of the independent associations, the leading SNP (the SNP that was most significant in any of the phenotypes or strata) was taken forward for replication. In total, 97 of the 133 independent associations (Supplementary Data 1) replicated (P<0.05/133). Out of these, 30 were associated with AFR, 42 with LFR and 65 with TFR in either the sex-combined or sex-stratified analyses. There was substantial overlap in associated loci between LFR and TFR loci (Figure 2) while associations with AFR overlapped to a small degree. One locus in the vicinity of *ADAMTSL3* was associated with all three phenotypes.

**Figure 2.**
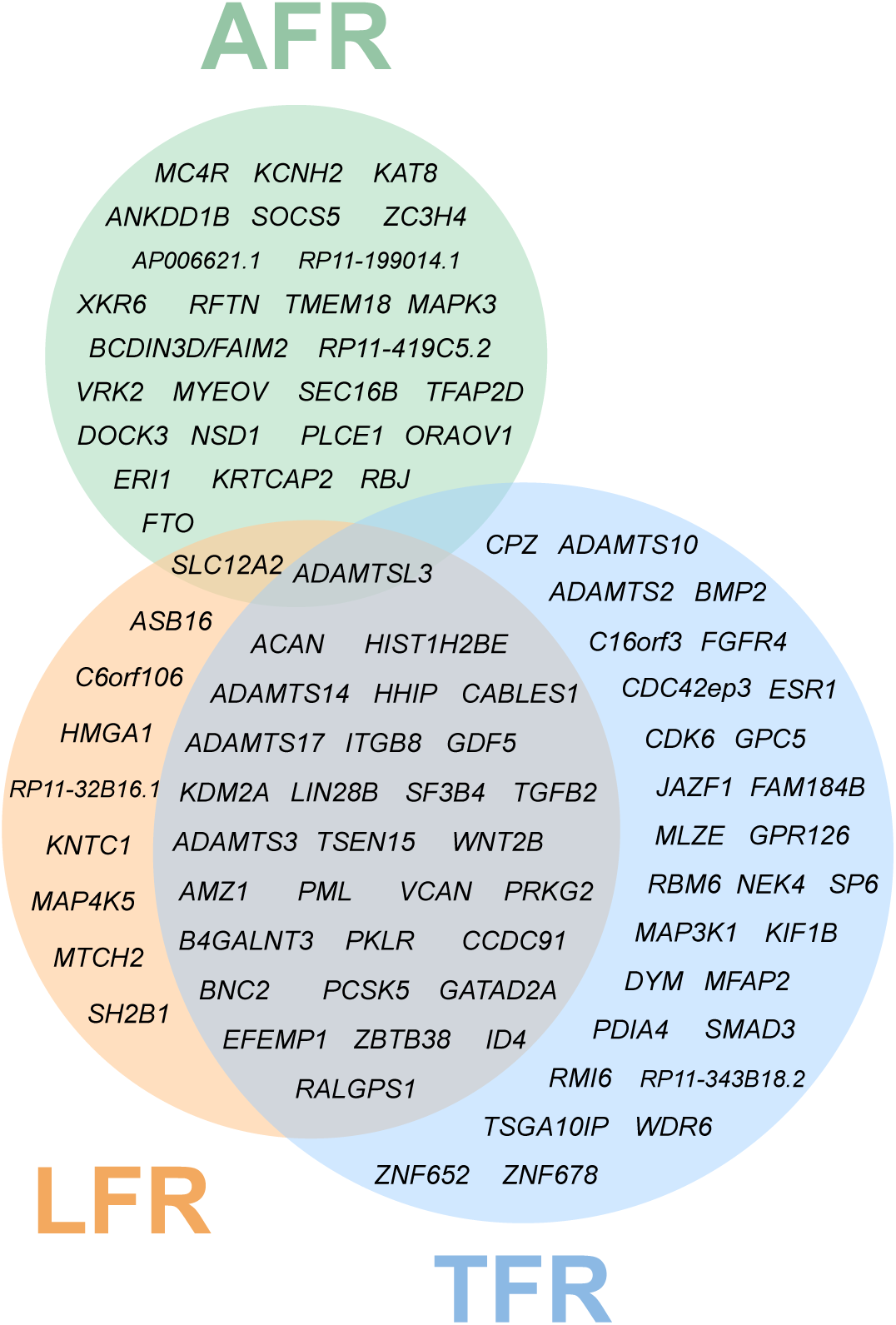
Body fat ratio associated loci and the overlap of associations between AFR, LFR and TFR. Loci are denoted by the nearest gene or by the most likely target gene (see methods section).

In the sex-stratified GWAS, only six loci were associated (replication P-value < 0.05/133) with body fat ratios in males while 71 loci were observed in females. Sex-heterogenous effects of associated variants were tested for using the GWAMA software. We observed 42 independent body fat ratio-associated variants whose effect differed between females and males (P < 5.15*10^−4^, Table 2). In all but two cases, the effects were stronger, or only present in females. Stronger effects in males were observed for two AFR-associated variants rs3812049 near *SLC12A2,* and rs11289753 near *PLCE1* (Table 2).

LD score regression (LDSC) was used to estimate the fraction of variance of body fat ratios that could be explained by SNPs, i.e. the SNP heritability^17^. SNP heritability was higher in females compared to males for all traits and ranged from ~20% to ~26% in females and from 12% to 16% in males (supplementary Table 2).

### Phenotypic and genetic correlation between body fat ratios and anthropometric phenotypes

Phenotypic correlations were assessed, in males and females separately, by calculating squared semi-partial correlation coefficients from the ANOVA table of two nested linear models that were adjusted for age and principal components, while genetic correlations were estimated using cross-trait LD score regression^19^ (see methods). Overall, the genetic and phenotypic correlations showed a large degree of similarity (supplementary Table 3 and 4) and the correlation between the anthropometric traits and the ratios were in the same direction for phenotypic and genetic correlations for all correlations that reached the threshold for significance. In females, BMI and WC was strongly correlated with AFR both with regards to phenotypic (BMI - 78.9% and WC - 54.3%; squared semi-partial correlation coefficients, see supplementary Table 3) and genetic (BMI - 79.2% and WC - 51,8%; squared genetic correlation coefficients, see supplementary Table 4) correlations. Height also contributed to a moderate degree in explaining the phenotypic variance in LFR and TFR in females (16.0% and 25.3%) even though the genotypic correlation between height and both LFR and TFR was even higher (42.3% and 60.8% variance explained). In males, anthropometric traits contributed only to a small degree, up to 4.8%, in explaining the phenotypic variance in the ratios. This is also supported by the lower genetic correlations between these traits with the highest correlation seen between BMI and TFR (14.4% variance explained).

There was also a strong correlation between LFR and TFR in both males and females (>82% of the variance explained in both males and females and for both genetic and phenotypic correlations, supplementary Table 3 and 4). LFR and TFR were inversely correlated, which agrees well with the large overlap in GWAS results for these phenotypes and the fact that the effect estimates from the GWAS was in the opposite direction for LFR and TFR (Supplementary Data 1). In contrast, AFR appeared to be more independent as only a low amount of phenotypic and genetic correlation was observed between AFR and the other two traits (supplementary Table 3 and 4).

### Overlap with findings from previous GWAS

Body fat ratio-associated SNPs were tested for overlap with associations from previous GWAS for anthropometric traits by determining LD with entries from GWAS-catalog^20^. We used a strict cut-off of *R^2^* < 0.1 to distinguish novel findings. In total, we identify 40 body fat ratio-associated signals that have not previously been associated with an anthropometric trait (Table 1). Forty body fat ratio associated signals overlapped with previously identified height-associated loci^21,22^ (Supplementary Data 1) and the majority of these signals were associated with TFR and/or LFR (36 out of 40). For AFR, the strongest associations were observed at well-known BMI and adiposity-associated loci such as: *FTO, MC4R, TMEM18, SEC16B* and *TFAP2B* (Supplementary Data 1), which agrees with the strong correlations (genotypic and phenotypic) between AFR and BMI.

**Table 1.**
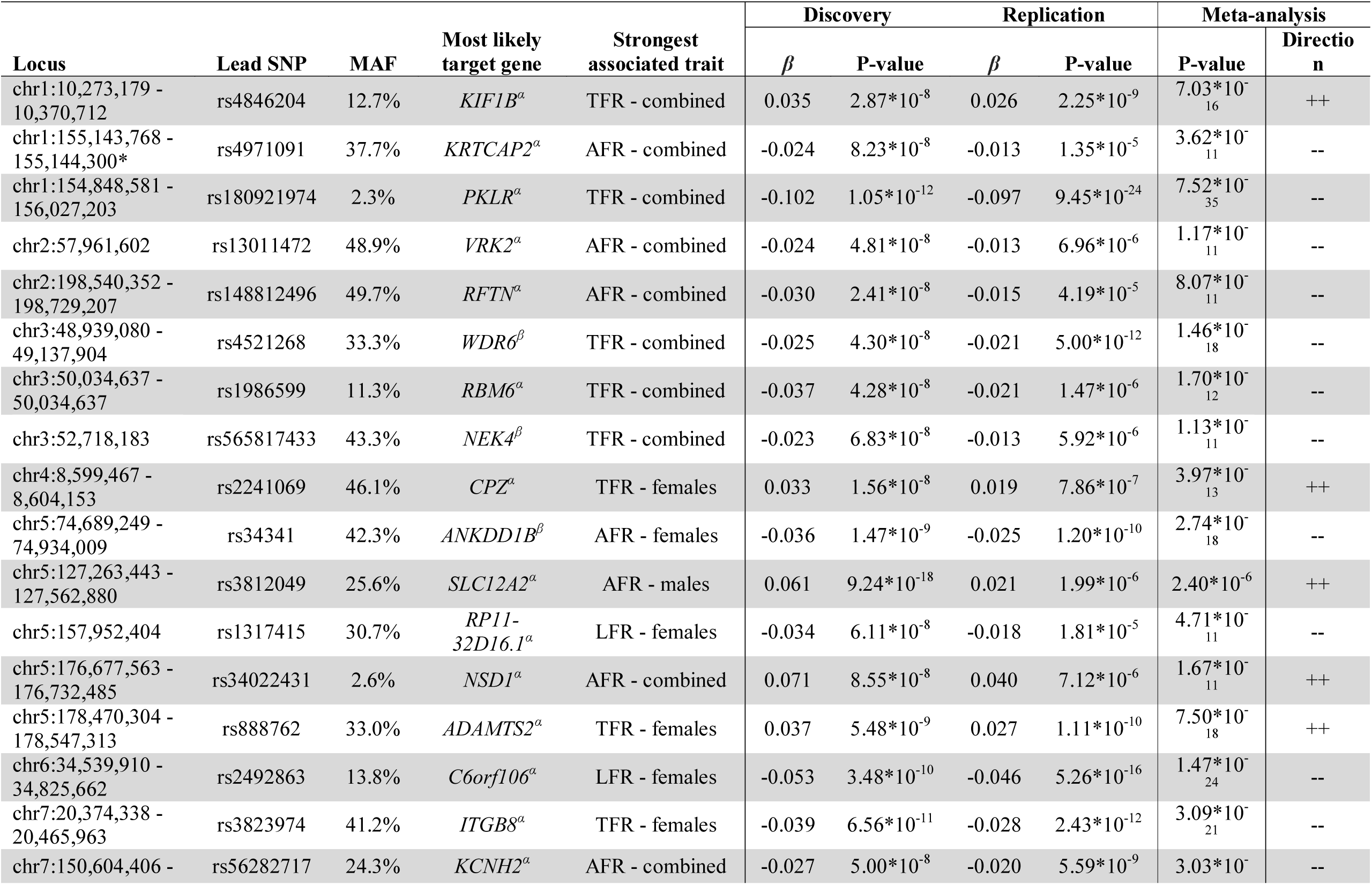
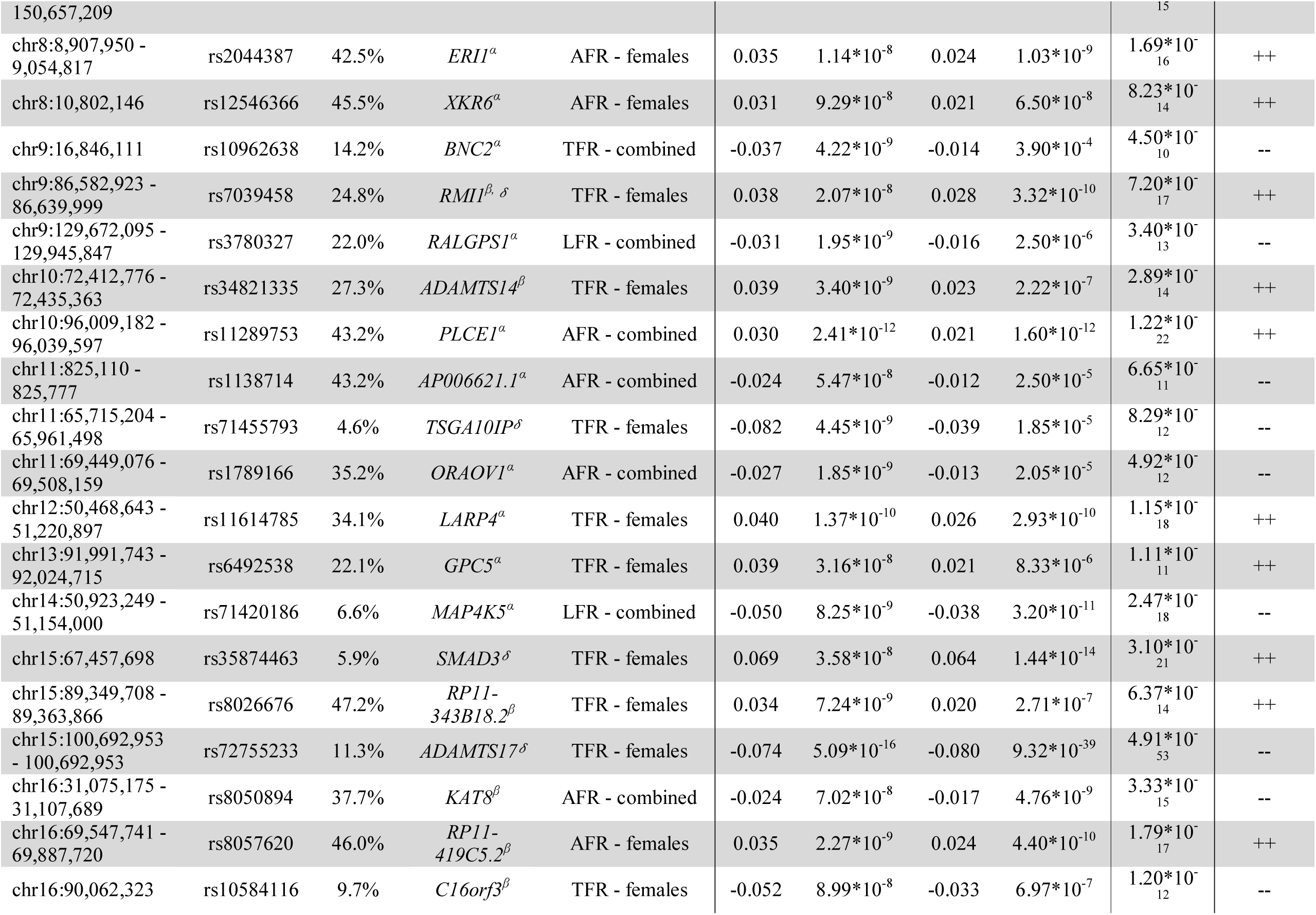
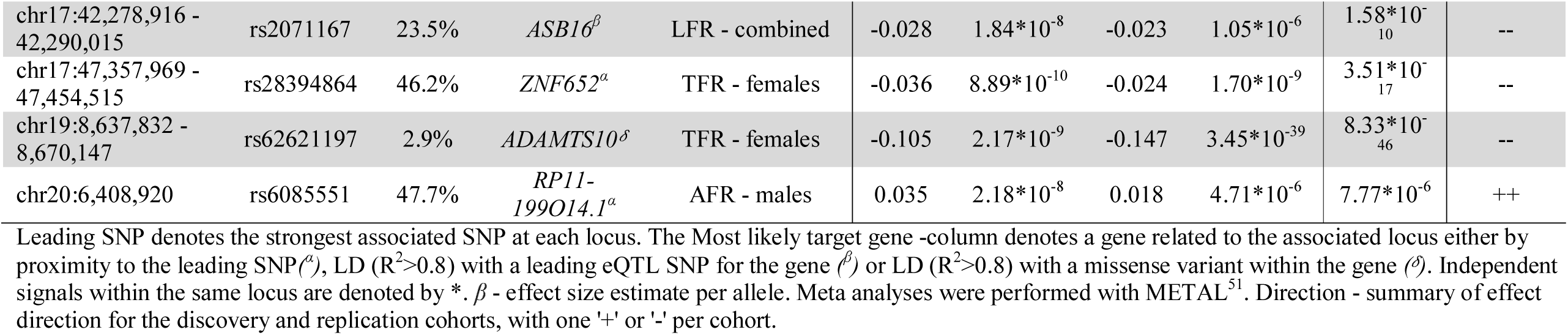
Novel body fat ratio-associated loci. Entries represent loci that have not previously been associated with an anthropometric trait.

We also compared the direction of the effects for overlapping GWAS results by estimating the effects of leading body fat ratio-associated SNPs on the respective overlapping anthropometric traits in the discovery cohort. The effects of TFR-associated SNPs were directionally consistent with effects on height and WHRadjBMI, while the effects were the opposite for LFR. The direction of effects for AFR-associated SNPs were consistent with effects on BMI-, WC- and WHR (supplementary Table 5), which agrees with the strong genetic and phenotypic correlation between AFR and these phenotypes.

Among the novel loci, five overlapped with traits associated with cardiovascular disease: two loci near *ANKDD1B* and *KAT8* that have been associated with LDL cholesterol^23^ and triglycerides^24^, respectively; and three loci near *KCNH2, XKR6* and *BMP2* that have been associated with QT interval^25-27^, thickness of the carotid intima media^28^ and QRS complex^29^, respectively (supplementary Table 6).

### Functional annotation of associated loci

Functional annotation of the GWAS loci was performed by identifying overlap with eQTLs from the Genotype-Tissue Expression (GTEx) project^30^ and by identifying potentially deleterious missense variants in LD (*R^2^* > 0.8) with our leading SNPs (see method section). In total, 31 body fat ratio-associated loci overlapped with an eQTL (Supplementary Data 2), and 11 leading SNPs were in LD with a potentially deleterious missense variant (Table 3). The probability for functional effects of missense variants has been predicted by sequence analyses^31,32^ and Polyphen and SIFT-scores were used to assess the deleteriousness of the variants. Plausible causal missense variants were found in *ACAN, ADAMTS17,* FGFR4 and *ADAMTS10,* where the leading SNPs were predicted to have functional effects (Table 3). Rs315855, within *FGFR4,* has also previously been shown to be associated with progression of cancer^33,34^ and to affect insulin secretion *in vitro*^35^.

**Table 2.**
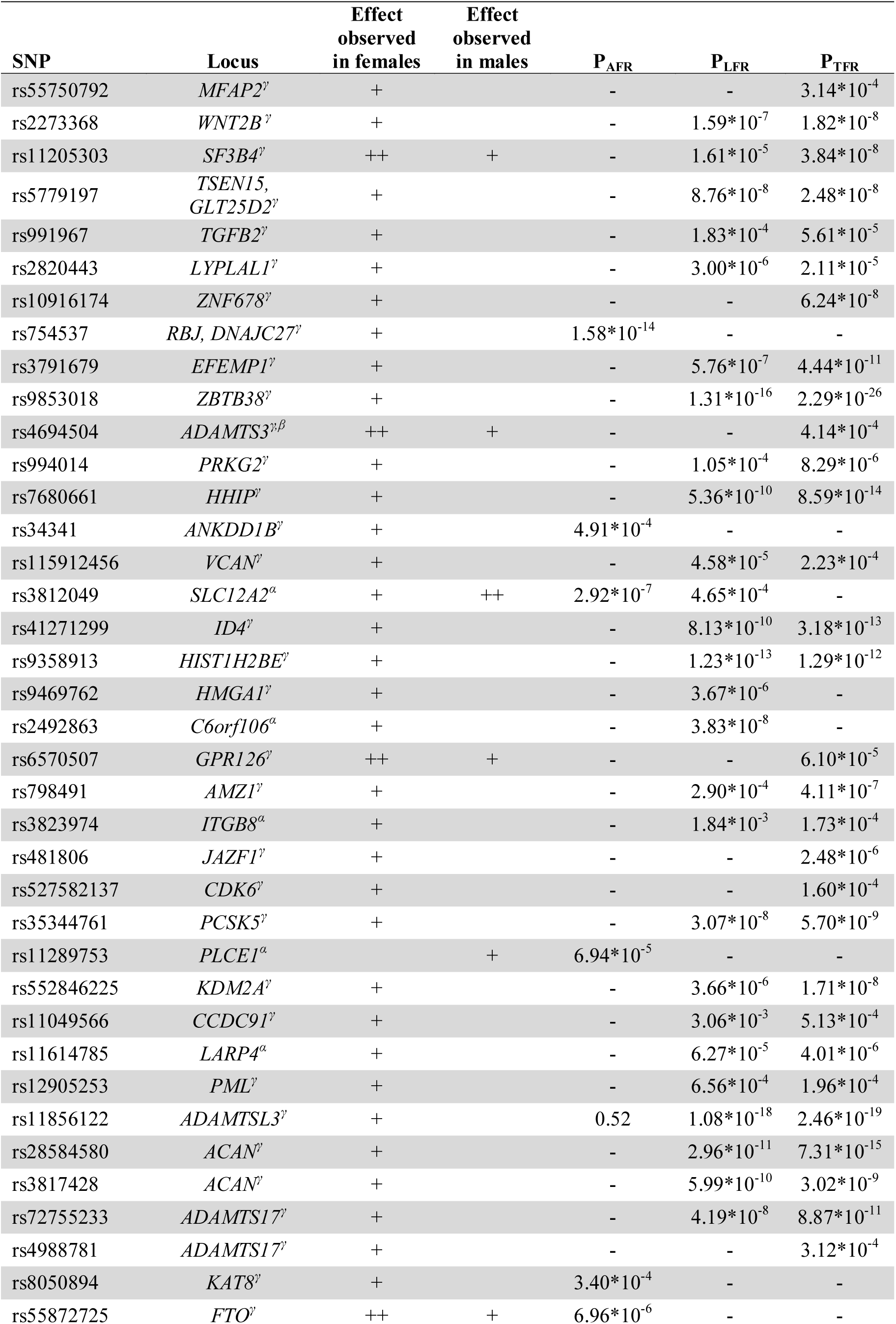
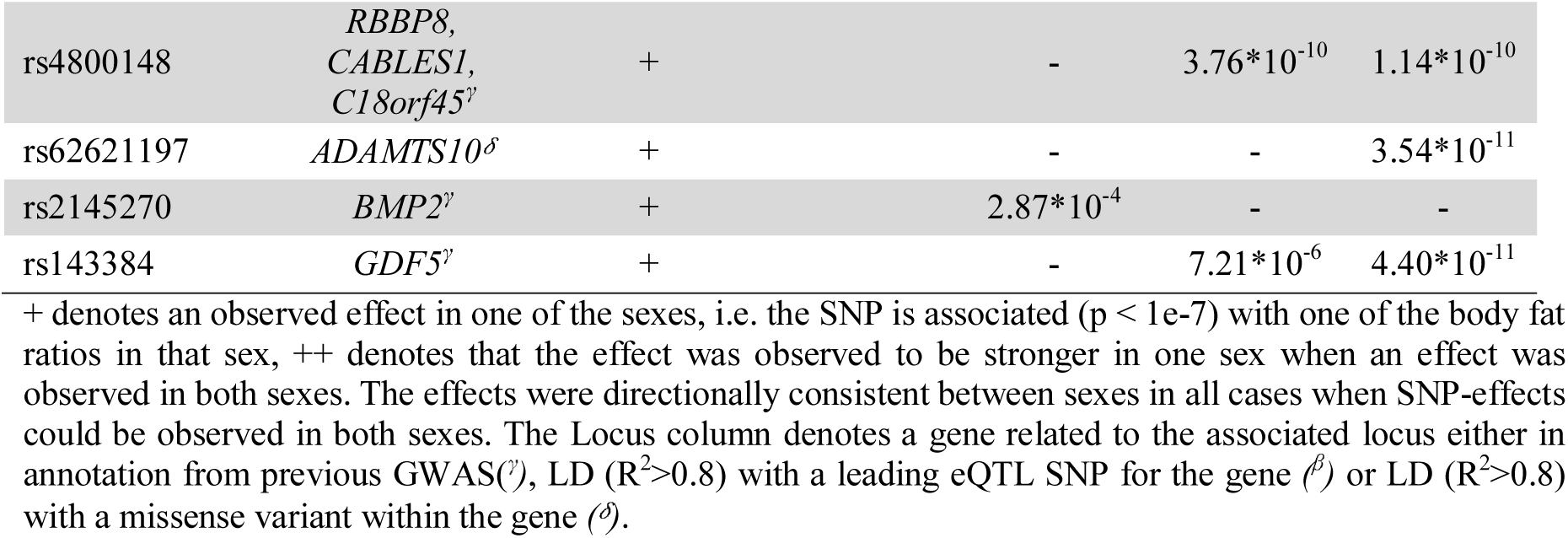
Sex heterogeneous effects of body fat ratio-associated SNPs was assessed with GWAMA^56^ for all replicated SNPs (N=97). P-values denote the results from tests for heterogeneity between sexes. Bonferroni correction was used to correct for multiple testing and P-values < 0.05/97 (5.15*10^−4^) were considered significant.

**Table 3.** Body fat ratio-associated potentially damaging missense variants.

| Leading SNP | Chr | Position (bp) | Missense variant | Position (bp) | Distance to leading SNP (bp) | LD w. leading SNP (R2) | Gene | Amino acid substitution | SIFT* score | Polyphen-2* score |
| --- | --- | --- | --- | --- | --- | --- | --- | --- | --- | --- |
| rs4733727 | 8 | 130,731,484 | rs4144738 | 130,760,850 | 29,366 | 0.91 | *GSDMC* | Met/Thr | deleterious | probably damaging |
| rs180921974 | 1 | 155,268,131 | rs72704117 | 155,175,089 | 93,042 | 0.95 | *THBS3* | Ala/Thr | deleterious | possibly damaging |
| rs3817428 | 15 | 89,415,247 | leading SNP |  |  |  | *ACAN* | Asp/Glu | deleterious | benign - possibly damaging |
| rs72755233 | 15 | 100,692,953 | leading SNP |  |  |  | *ADAMTS17* | Thr/Ile | deleterious | benign |
| rs115912456 | 5 | 82,815,158 | rs61749613 | 82,815,170 | 12 | 1.00 | *VCAN* | Lys/Glu | tolerated - deleterious | possibly damaging |
| rs9358913 | 6 | 26,239,404 | rs7766641 | 26,184,102 | 55,302 | 0.82 | *HIST1H2BE* | Gly/Ser | tolerated | probably damaging |
| rs351855 | 5 | 176,520,243 | leading SNP |  |  |  | *FGFR4* | Gly/Arg | tolerated | possibly damaging |
| rs71420186 | 14 | 50,960,918 | rs12881869 | 50,923,249 | 37,669 | 0.96 | *MAP4K5* | Ala/Thr | tolerated | possibly damaging |
| rs5779197 | 1 | 184,009,826 | rs2274432 | 184,020,945 | 11,119 | 0.96 | *TSEN15* | Gly/Asp | tolerated | benign - possibly damaging |
| rs143384 | 20 | 34,025,756 | rs224331 | 34,022,387 | 3,369 | 0.73 | *GDF5;*<br>*GDF5OS* | Ala/Ser;<br>Ala/Glu | tolerated | benign - possibly damaging |
| rs62621197 | 19 | 8,670,147 | leading SNP |  |  |  | *ADAMTS10* | Arg/Gln | tolerated | benign - possibly damaging |

\*Polyphen-2<sup>32</sup> and SIFT<sup>31</sup> scores were extracted from Ensembl (www.ensembl.org). SIFT utilizes protein sequence data to predict the effects of a missense variant while Polyphen-2 also considers protein structure and evolutionary conservation. Polyphen-2 generates a probability that a missense mutation is damaging, while SIFT generates a probability that the amino acid change is tolerated. Polyphen-2 uses a tree-tiered prediction score: 0-0.15 is predicted to be benign, 0.15 to 0.85 possibly damaging and 0.85 -1.0 more confidently predicted to be damaging (i.e., probably damaging). A SIFT score < 0.05 is predicted to be deleterious<sup>31</sup>. Polyphen-2 and SIFT scores are presented as ranges in the cases where they differ due to being part of different transcript variants.

### Enrichment analyses

To identify the functional roles and tissue specificity of associated variants, we performed enrichment analyses with DEPICT (Data-driven Expression Prioritized Integration for Complex Traits^36^, see method section). In these analyses we used summary statistics from sex-stratified GWAS on the combined cohort (195,043 women and 167,408 men) in order to maximize statistical power. Enrichment was only detected for TFR- and LFR-associated genes in females as well as LFR-associated genes in males (supplementary Data 3). We identified 212 enriched gene sets and there was a substantial overlap of enriched gene sets between TFR- and LFR- in females as well as moderate overlap with LFR-associated gene sets in males (supplementary Figure 5). The large fraction of overlapping gene sets between LFR and TFR in females agrees well with the large overlap in GWAS signals. Gene sets related to bone morphology and skeletal development (abnormal skeleton morphology, short limbs, decreased length of long bones, skeletal system morphogenesis) were among the most strongly associated with both LFR and TFR. We also find the ‘TGFβ signaling pathway’ gene set to be enriched for genes within the TFR and LFR-associated loci in females, as well as TGFβ downstream mediators: ‘SMAD1-’, ‘SMAD2-’, ‘SMAD3-’ and ‘SMAD7 protein-protein interaction subnetworks’ (supplementary Data 3).

The results from enrichment analyses were compared with results from previous GWAS for height^21^, BMI^9^ and WHRadjBMI^12^. Gene-sets that were enriched for LFR and TFR-associated genes were compared to results from previous GWAS for height^21^, BMI^9^ and WHRadjBMI^12^. Substantial overlap of enriched gene sets was observed between LFR/TFR-with height and WHRadjBMI (Figure 3a). In contrast, BMI associated biological processes overlapped only to a marginal extent: 2% of all gene sets that enriched for BMI-associated genes were also enriched for LFR- and TFR-associated genes.

**Figure 3.**
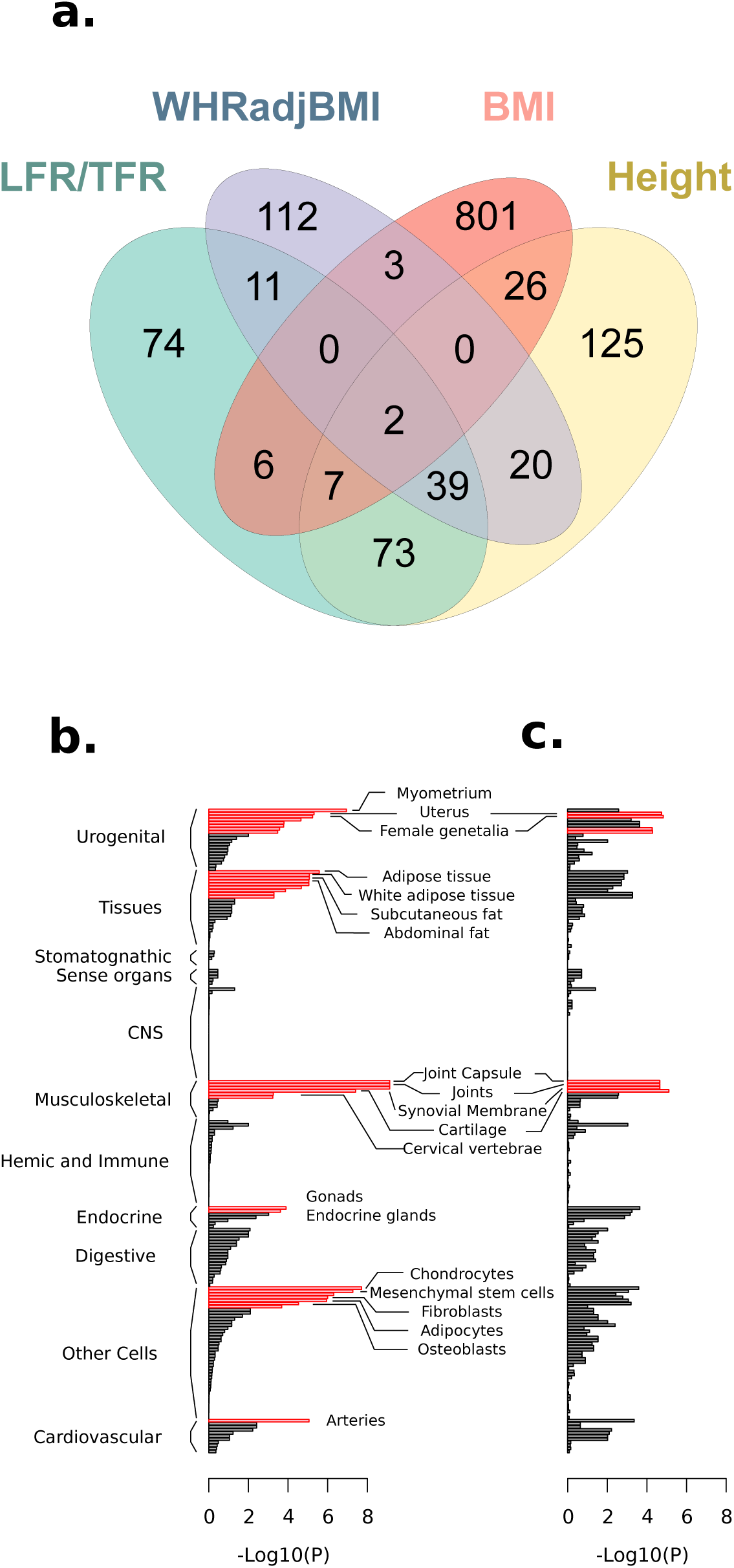
Enrichment analyses of genes at LFR and TFR associated loci. (**a**) Reconstituted gene sets found to be enriched for TFR- and LFR-associated genes (in males and females) were compared with results from previous GWAS on WHRadjBMI^12^, BMI^9^ and height^55^. Tissue and cell type enrichment of (**b**) TFR- and (**c**) LFR-associated genes. Red bars denote tissues gene sets that were significantly enriched for LFR- and TFR-associated genes at FDR < 0.05/12.

Tissue enrichment was observed for LFR and TFR-associated genes in females (Figure 3b-c) in gene sets related to female reproduction, musculoskeletal systems, chondrocytes, mesenchymal stem cells, and fibroblasts. For TFR, DEPICT also revealed enrichment of genes associated with adipose tissue cells as well as endocrine and cardiovascular systems (Figure 3). Tissue enrichment was not seen for the other traits or strata.

## Discussion

In this study, we performed GWAS on distribution of body fat to different compartments of the human body and identified and replicated 97 independent associations of which 40 have not been associated with any adiposity related phenotype previously. In contrast to previous studies, we have not addressed the total amount of fat but rather the fraction of the total body fat mass that is located in the arms, legs and trunk. Body fat distribution is well-known to differ between males and females, which we also clearly show in our study. We also show that the genetic effects that influence fat distribution are stronger in females compared to males. These results are consistent with previous GWAS that have revealed sexual dimorphisms in genetic loci for adiposity-related phenotypes, such as waist-circumference and waist-to-hip ratio^10,37,38^. Phenotypic and genetic correlations, as well as results from GWAS and subsequent enrichment analyses, also revealed that the amount of fat stored in the arms in females is highly correlated with BMI and WC. This suggests that the proportion of fat stored in the arms will generally increase with increased accumulation of body mass and adipose tissue. In contrast, males exhibited moderate- to weak phenotypic and genetic correlations between the distribution of fat to different parts of the body and anthropometric traits, which indicates that the proportions of body fat mass in different compartments of the male body remains more stable as body mass and body adiposity increases. Among the three phenotypes analyzed in this study LFR and TFR were inversely correlated in both males and females. This suggests that LFR and TFR to a large extent describe one trait, i.e. the distribution of adipose tissue between these two compartments which is further supported by the large overlap in GWAS loci between the two phenotypes. In contrast, AFR was only weakly correlated with the other two traits.

Tissue enrichment revealed an important role in body fat distribution in females for mesenchyme derived tissues: i.e. adipose and musculoskeletal tissues; as well as tissues related to female reproduction. This suggests that the distribution of fat to the legs and trunk in females is mainly driven by the effect of female gonadal hormones on mesenchymal progenitors of musculoskeletal and adipose tissues. Enrichment analyses also showed that LFR and TFR have unique features that separates them from other anthropometric measurements, which was indicated by the portion of LFR/TFR-associated gene sets that did not overlap with height-, BMI or WHRadjBMI-associated gene sets. However, there was also an overlap in the functional aspects between these traits with both height and WHRadjBMI. This is indicated by the tissue enrichment profile for LFR/TFR-associated genes, which shares features with tissue enrichments reported for height in previous GWAS^39^: height-associated genes were strongly enriched in musculoskeletal tissue types with additional enrichment in cardiovascular and endocrine tissue types; as well as WHRadjBMI^12^: associated genes were enriched in adipocytes and adipose tissue subtypes. Of particular note, we did not identify any enrichment of body fat ratio-associated genes in CNS tissue gene sets in contrasts to enrichment analyses in previous GWAS for BMI where CNS have been indicated to play a prominent role in obesity susceptibility^9^.

In the GWAS for LFR and TFR in females, we find that several genes that highlight the influence of biological processes related to the interaction between cells and the extracellular matrix (ECM), as well as ECM-maintenance and remodeling. These include *ADAMTS2, ADAMTS3, ADAMTS10, ADAMTS14,* and *ADAMTS17,* which encode extracellular proteases that are involved in enzymatic remodeling of the ECM. Two leading SNPs were in LD with potentially damaging missense mutations in *ADAMTS10* and *ADAMTS17* and two other GWAS signals overlapped with eQTLs for *ADAMTS14* and *ADAMTS3.* In addition, possibly deleterious missense mutations in LD with our leading GWAS SNPs were also found for *VCAN* and *ACAN.* Both *VCAN* and *ACAN* encode chondroitin sulfate proteoglycan core proteins that constitute structural components of the extracellular matrix, particularly in soft tissues^40^. These proteins also serve as major substrates for ADAMTS proteinases^41^. ECM forms the three-dimensional support structure for connective and soft tissue. In fat tissue, the ECM regulates adipocyte expansion and proliferation^42^. Remodeling of the ECM is required to allow for adipose tissue growth and this is achieved through enzymatic processing of extracellular molecules such as proteoglycans, collagen and hyaluronic acid. For example, ADAMTS2, 3 and 14 act as procollagen N-propeptidases that mediate the maturation of triple helical collagen fibrils^43,44^. We therefore propose that effects of genetic variation in biological systems involved in ECM-remodeling as a factor underlying normal variation in female body fat distribution.

## Conclusions

In summary, GWAS of body fat distribution determined by sBIA reveals a genetic architecture that influences distribution of adipose tissue to the arms, legs and trunk. Genetic associations and effects clearly differ between sexes, in particular for distribution of adipose tissue to the legs and trunk. The distribution of body fat in women has been previously been suggested as a causal factor leading to lower risk of cardiovascular and metabolic disease, as well as cardiovascular mortality for women in middle age^5^ and genetic studies have identified SNPs that are associated with a ‘favorable’ body fat distribution^45^, i.e. with higher BMI but lower risk of cardiovascular and metabolic disease. The capacity for peripheral adipose storage has been highlighted as one of the components underlying this phenomenon^45^. Resolving the genetic determinants and mechanisms that lead to a favorable distribution of body fat may help in risk assessment and in identifying novel venues for intervention to prevent or treat obesity-related disease.

## Author contributions

MRA, TK, WE and ÅJ conceived of and designed the study. Analysis was performed by MRA and WE under supervision by ÅJ. MRA analyzed the data and wrote the first draft of the manuscript. All authors contributed to the final version of the manuscript.

## Acknowledgements

We are grateful to the participants and staff of the UK Biobank. Access to UK Biobank genetic and phenotypic data was granted under application no. 15152. Computations were performed on the computational cluster at the Uppsala Multidisciplinary Center for Advanced Computational Science (UPPMAX) under projects b2016021 and b2017066. The work was supported by grants from the Swedish Society for Medical Research (SSMF), the Kjell and Märta Beijers Foundation, Göran Gustafssons Foundation, the Swedish Medical Research Council (Project Number 2015-03327), the Marcus Borgström Foundation, and the Åke Wiberg Foundation.

## online Methods

### UK Biobank participants

The first release of imputed genotype data from UK Biobank (N = 152,249) was used as a discovery cohort, and imputed genotype data from an unrelated set of participants from the third genotype release (N = 326,565) as a replication cohort. Participants who self-reported as being of British descent (data field 21000) and were classified as Caucasian by principal component analysis (data field 22006) were included in the analysis. Genetic relatedness pairing was provided by the UK Biobank (Data field 22011). Participants were removed due to relatedness based on kinship data (estimated genetic relationship > 0.044), poor genotyping call rate (<95%), high heterozygosity (Data field 22010), or sex-errors (Data filed 22001). After filtering, 116,138 participants remained in the discovery cohort and 246,361 in the replication cohort and were included in downstream analyses.

### Ethics

Ethical approval to collect participant data was given by the North West Multicentre Research Ethics Committee, the National Information Governance Board for Health & Social Care, and the Community Health Index Advisory Group. UK Biobank possesses a generic Research Tissue Bank approval granted by the National Research Ethics Service (http://www.hra.nhs.uk/), which lets applicants conduct research on UK Biobank data without obtaining separate ethical approvals. Access to UK Biobank genetic and phenotypic data was granted under application no. 15152. All participants provided signed consent to participate in UK Biobank ^46^.

### Genotyping, imputations and QC

Genotyping in the discovery cohort had been performed on two custom-designed microarrays: referred to as UK BiLEVE and Axiom arrays, which genotyped 807,411 and 820,967 SNPs, respectively. Imputation had been performed using UK10K^47^ and 1000 genomes phase 3^48^ as reference panels. Prior to analysis, we filtered SNPs based on call rate (--geno 0.05), HWE (P-value > 10^−20^, MAF (--maf 0.0001) and imputation quality (Info > 0.3) resulting in 25,472,837 SNPs in the discovery cohort. The third release of data from the UK Biobank contained genotyped and imputed data for 488,366 participants (partly overlapping with the first release). For our replication analyses, we included an independent subset that did not overlap with the discovery cohort. Genotyping in this subset was performed exclusively on the UK Biobank Axiom Array. This dataset included 47,512,111 SNPs that were filtered based on HWE (P<10^−20^), call rate (--geno 0.05), (Info >0.3) and MAF (--maf 0.0001). All genomic positions are in reference to hg19/build 37.

### Phenotypic measurements

The phenotypes used in this study derive from impedance measurements produced by the Tanita BC418MA body composition analyzer. Participants were barefoot, wearing light indoor clothing, and measurements were taken with participants in the standing position. Height and weight were entered manually into the analyzer before measurement. The Tanita BC418MA uses eight electrodes: two for each foot and two for each hand. This allows for five impedance measurements: whole body, right leg, left leg, right arm and left arm (Figure 1a.). Body fat for the whole body and individual body parts had been calculated using a regression formula, that was derived from reference measurements of body composition by DXA (Figure 1b) in Japanese and Western subjects. This formula uses weight, age, height and impedance measurements^49^ as input data. Arm, and leg fat masses were averaged over both limbs. Arm, leg, and trunk fat masses were then divided by the total body fat mass to obtain the ratios of fat mass for the arms, legs and trunk, i.e. what proportion of the total fat in the body is distributed to each of these compartments. These variables were analyzed in this study and were named: arm fat ratio (AFR), leg fat ratio (LFR), and trunk fat ratio (TFR).

### Assessing the relationship between adipose tissue ratios and anthropometric traits

Phenotypic correlations between fat distribution ratios and anthropometric traits were estimated by calculating semi-partial correlation coefficients for males and females separately, using anova.glm in R. Adipose tissue ratios (AFR, LFR or TFR) were set as the response variable. BMI, waist circumference, waist circumference adjusted for BMI, waist-to-hip ratio, height, or one of the other ratios were included as the last term in a linear model that included, age and principal components as covariates. The reduction in residual deviance, i.e., the reductions in the residual sum of squares as BMI, waist circumference, waist circumference adjusted for BMI, waist-to-hip ratio, height, or one of the other ratios was added to the model, is presented as percentages of the total deviance of the null model in supplementary Table 3.

### Genome wide association study for body fat ratio

A two-stage GWAS was performed using a discovery and a replication cohort. Body fat ratios were adjusted for age, age squared and normalized by rank-transformation separately in males and females using the *rntransform* function included in the GenABEL library^50^. GWAS was performed in PLINK v1.90b3n^18^ using linear regression models with AFR, LFR, and TFR as the response variables and the SNPs as predictor variables. A batch variable was used as covariate in the GWAS for the discovery analyses to adjust for genotyping array (Axiom and BiLEVE). We also included age, the first 15 principal components and sex (in the sex-combined analyses) as covariates in the GWAS. LD score regression intercepts (see further information below), calculated using ldsc^17^, were used to adjust for genomic inflation, by dividing the square of the t-statistic for each tested SNP with the LD-score regression intercept for that GWAS, and then calculating new p-values based on the adjusted t-statistic. We used a threshold of P<10^−7^, after adjusting for LD score intercept, as threshold for significance in the discovery cohort.

The -clump function in PLINK was used to identify the number of independent signals in each GWAS. This function groups associated SNPs based on the linkage disequilibrium (LD) pattern. The parameters for clumps were set to: -clump-p1 1*10^−7^, clump-p2 1*10^−7^, clump-r2 0.10 and --clump-kb 1000. This function groups SNPs within one million base pairs that were associated with the trait at p < 1*10^−7^. Several associations were found in more than one of the three body fat ratios (AFR, TFR, or LFR) or strata (males, females or sex-combined) and different leading SNPs were observed for different traits and strata at several loci. To assess whether these represented the same signal, we assessed the LD between overlapping leading SNPs in PLINK. SNPs in low LD (*R^2^*-value < 0.05) were considered to represent independent signals. For leading SNPs where we could not exclude that they were in LD (*R^2^-*value >= 0.05) we performed conditional analysis in PLINK, conditioning on the most significant SNP across all phenotypes and strata. Associations with a P< 1*10^−7^ after conditioning for the most significant SNP were considered as being independent signals. For each independent signal, the leading SNP (lowest p-value) was taken forward for replication. Meta analyses of results from the discovery and replication cohorts was performed with the METAL software^51^ for all independent associations that were taken forward for replication.

### SNP heritability, and genetic correlations

We estimated SNP heritability and genetic correlations using LD score regression (LDSC), implemented in the ‘ldsc’ software package^17^. LDSC uses LD patterns and summary stats from GWAS as input. For genetic correlations, we performed additional sex-stratified GWAS in the UK biobank (using the same covariates as for the ratios) for standard anthropometric traits, BMI, height, WC, WHR, WCadjBMI and WHRadjBMI, in the discovery cohort. GWAS summary stats were filtered for SNPs included in HapMap3 to reduce likelihood of bias induced by poor imputation quality. After this filtering, 1,164,192 SNPs remained for LDSC analyses. LD scores from the European data of the 1000 Genomes project (including LD patterns for all the HapMap3 SNPs) for use with LDSC were downloaded from the Broad institute at: https://data.broadinstitute.org/alkesgroup/LDSCORE/eur_w_ld_chr.tar.bz2. Genetic correlations between the three body fat ratios and anthropometric traits were assessed by cross-trait LD score regression.

### Overlap with findings from previous GWAS

Leading SNPs from all independent signals in our analyses were cross referenced with the NHGRI-EBI catalog of published genome-wide association studies (GWAS Catalog - data downloaded on 23 April 2018) ^20^ to determine if body fat ratio-associated signals overlapped with previously identified anthropometric associations from previous GWAS. We used a cutoff of *R^2^* < 0.1 between SNPs from our analyses and anthropometric trait-associated SNPs (P< 5*10^−8^) from GWAS catalog to determine if an association was novel. LD between data in the GWAS catalogue and our leading SNPs were calculated using PLINK v1.90b3n^18^. In addition, leading SNPs at Body fat ratio-associated loci that potentially overlapped *(R^2^* > 0.1) with signals from previous GWAS were tested for association with standard anthropometric traits (BMI, height, WC, WHR, WCadjBMI and WHRadjBMI) in the UK biobank discovery cohort using PLINK v1.90b3n^18^ through linear regression modelling and including sex, age a batch variable and 15 principal components as covariates. Here, a P-value of P < 1e-7 was considered significant.

### Functional annotation of associated loci

Associated loci were investigated for overlap with eQTLs from the GTEx project ^30^. The threshold for significance for the eQTLs was set to 2.3*10^−9^ in agreement with previous studies^52^. The strongest associated SNP for each tissue and gene in the GTEx dataset was identified. We then estimated the LD between the top eQTL SNPs and the leading SNPs for each independent association from our analysis. If a SNPs from our analyses were in LD *(R^2^* > 0.8) with a leading eQTL SNP the two signals were considered overlapping.

Leading SNPs, and all SNPs in LD *(R^2^* > 0.8) with a leading SNPs from our analyses (LD determined in the UK biobank cohort in PLINK) were cross-referenced with dbSNP (human 9606 b150) in order to identify potentially deleterious intragenic variants in LD (*R*^2^ > 0.8) with body fat ratio-associated variants. Polyphen and SIFT-scores for the missense variants (extracted from Ensembl - www.ensembl.org) were used to assess the deleteriousness of the variants.

### Enrichment analysis

To identify the functional roles and tissue specificity of associated variants, we performed tissue and gene-set enrichment analyses using DEPICT^36^. For the gene-set enrichment in DEPICT, gene expression data from 77,840 samples have been used to predict gene function for all genes in the genome based on similarities in gene expression. In comparison to standard enrichment tools that apply a binary definition to define membership in a set of genes that have been associated with a biological pathway or functional category (genes are either included or not included), in DEPICT, the probability of a gene being a member of a gene set has instead been estimated based on correlation in gene expression. This membership probability to each gene set has been estimated for all genes in the human genome and the membership probabilities for each gene have been designated ‘reconstituted’ gene sets. A total of 14,461 reconstituted gene sets have been generated which represent a wide set of biological annotations (Gene Ontology [GO], KEGG, REACTOME, Mammalian Phenotype [MP], etc.). For tissue enrichment in DEPCIT, microarray data from 37,427 human tissues have been used to identify genes with high expression in different cells and tissues. When performing the tissue enrichment analyses in DEPICT, the most significant GWAS hits are scanned for enrichment of genes that are highly expressed in any cells or tissues.

For the enrichment analyses, we performed sex-stratified GWAS for AFR, LFR and TFR on a combined set of the UK Biobank cohort, in order to achieve high power for enrichment analyses, including 195,043 females and 167,408 males. The ‘clump’ functionality in PLINK is used to determine associated loci. The p-value cut off for association for ‘clump’ was set at P < 10^−7^. In the enrichment analyses, DEPICT assesses whether the reconstituted gene sets are enriched for genes within trait-associated loci^36^. The False Discovery Rate (FDR)^53^ was used to adjust for multiple testing. Twelve analyses were run in total (tissue enrichment and gene-set enrichment) and FDR<0.05/12 was considered significant.

### Interaction between SNPs and sex

We used the GWAMA software^54^ to test for heterogenous effects of associated SNPs between sexes. In GWAMA, fixed-effect estimates of sex-specific and sex-combined beta coefficients and standard errors are calculated from GWAS summary statistics to test for heterogeneous allelic effects between females and males. GWAMA obtains a test-statistic by subtracting the sex-combined squared t-statistic from the sum of the two sex-specific squared t-statistics. This test statistic is asymptotically chi-square-distributed and equivalent to a normal z-test of the difference in allelic effects between sexes. SNPs that were significant in the replication analyses in any of the strata were tested for heterogeneity between sexes in the replication cohort.

Bonferroni correction was used to correct for multiple testing and p-values < 0.05/97 were considered to be significant.

### Data availability

The data that support the findings of this study are available from UK Biobank (http://www.ukbiobank.ac.uk/about-biobank-uk/). Restrictions apply to the availability of these data, which were used under license for the current study (project no. 15152). Data is available for bona fide researchers upon application to the UK Biobank. Summary statistics from all association tests are available for download at: https://myfiles.uu.se/ssf/s/readFile/share/3993/1270878243748486898/publicLink/GWAS_summary_stats_ratios.zip.

## Supplementary Figure and Tables

**Supplementary Table 1. Basic characteristics of UK Biobank participants included in the analyses.**

**Supplementary Table 2. SNP-heritability *(h2)* and LD score regression intercept estimates of body fat ratios.**

**Supplementary Table 3. Phenotypic correlation between body fat ratios and anthropometric traits.**

**Supplementary Table 4. Genetic correlations between body fat ratios and anthropometric traits.**

**Supplementary Table 5. Direction of effect for body fat ratio-associated leading SNPs for standard anthropometric traits.**

**Supplementary Table 6. Overlap between body fat ratio-associated loci with non-anthropometric trait-associations from previous GWAS.**

**Supplementary Figure 1. Quantile-quantile (QQ) plots of the distribution of p-values from GWAS of body fat ratios in the UK Bibank sex-combined, female and male discovery cohorts.**

**Supplementary Figure 2. Manhattan plots for GWAS results for arm fat ratio in the combined discovery cohort and sex-stratified analyses.**

**Supplementary Figure 3. Manhattan plots for GWAS results for leg fat ratio in the combined discovery cohort and sex-stratified analyses.**

**Supplementary Figure 4. Manhattan plots for GWAS results for trunk fat ratio in the combined discovery cohort and sex-stratified analyses.**

**Supplementary Figure 5. Overlap of enriched gene sets from DEPICT for body fat ratios.**

## Supplementary data sets

**Supplementary Data 1. Independent signals for association with body fat ratios.**

**Supplementary Data 2. Expression quantitative loci (eQTLS) in LD with leading SNPs at body fat ratio-associated loci.**

**Supplementary Data Set 3. Results from enrichment analyses by DEPICT.**

